# Comparison of three TaqMan Real-Time Reverse Transcription-PCR assays in detecting SARS-CoV-2

**DOI:** 10.1101/2020.07.06.189860

**Authors:** Yan Xiao, Zhen Li, Xinming Wang, Yingying Wang, Ying Wang, Geng Wang, Lili Ren, Jianguo Li

## Abstract

Quick and accurate detection of SARS-CoV-2 is critical for COVID-19 control. Dozens of real-time reverse transcription PCR (qRT-PCR) assays have been developed to meet the urgent need of COVID-19 control. However, methodological comparisons among the developed qRT-PCR assays are limited. In the present study, we evaluated the sensitivity, specificity, amplification efficiency, and linear detection ranges of three qRT-PCR assays, including the assays developed by our group (IPBCAMS), and the assays recommended by WHO and China CDC (CCDC). The three qRT-PCR assays exhibited similar sensitivities, with the limit of detection (LOD) at about 10 copies per reaction (except the ORF 1b gene assay in CCDC assays with a LOD at about 100 copies per reaction). No cross reaction with other respiratory viruses were observed in all of the three qRT-PCR assays. Wide linear detection ranges from 10^6^ to 10^1^ copies per reaction and acceptable reproducibility were obtained. By using 25 clinical specimens, the N gene assay of IPBCAMS assays and CCDC assays performed better (with detection rates of 92% and 100%, respectively) than that of the WHO assays (with a detection rate of 60%), and the ORF 1b gene assay in IPBCAMS assays performed better (with a detection rate of 64%) than those of the WHO assays and the CCDC assays (with detection rates of 48% and 20%, respectively). In conclusion, the N gene assays of CCDC assays and IPBCAMS assays and the ORF 1b gene assay of IPBCAMS assays were recommended for qRT-PCR screening of SARS-CoV-2.

## Introduction

Since the first detection in late 2019, severe respiratory syndrome CoV-2 (SARS-CoV-2) caused Corona Virus Infectious Disease in 2019 (COVID-19) has widely spread in the world. By April 11, 2020, more than 1.7 million patients infected by SARS-CoV-2 has been reported from 185 countries (1). Given the quick increase in confirmed cases and asymptomatic infections, there are increasing demands in diagnostic tools for quick and accurate detection of the virus (2, 3). Several real-time reverse transcription-Polymerase Chain Reaction (qRT-PCR) for the detection of SARS-COV-2 has been developed to meet the demands, including the assays by this group (IPBCAMS assays), and the assays by WHO (WHO assays), and the assays by China CDC (CCDC assays).

Because SARS-CoV-2 usually infected the lower respiratory tract, it is not easy to detect the viral nucleic acids from throat swabs with relatively lower viral load (4). Thus, qRT-PCR assays with higher sensitivity and better performance in the detection of SARS-CoV-2 is recommended in aiding the diagnosis of COVID-19 (2). However, most of the current available qRT-PCR assays were developed for emergency, a comprehensive methodological comparison among these assays remains unfulfilled.

To comprehensively compare the performance of currently available qRT-PCR assays for detection of SARS-CoV-2, we evaluated the sensitivity, specificity, amplification efficiency, and linear detection ranges among IPBCAMS assays, WHO assays and CCDC assays.

## Materials and methods

### Nucleic acid extraction

Nucleic acids were extracted from a volume of 200 μl clinical samples by using NucliSens easyMag apparatus (bioMerieux, MarcyL’Etoile, France) according to the manufacturer’s instructions. A volume of 50 μl total nucleic acid eluate for each specimen was recovered and transferred into a nuclease-free vial and either tested immediately or stored at −80°C.

### Primers and probes

Sequences of primers and probes for the IPBCAMS assays were recently developed (5), while those for the WHO assays were obtained from the website of WHO (https://www.who.int/docs/default-source/coronaviruse/protocol-v2-1.pdf?sfvrsn=a9ef618c_2), and those for the CCDC assays were obtained from the website of China CDC (http://www.chinacdc.cn/jkzt/crb/zl/szkb_11803/jszl_11815/202003/W020200309540843062947.pdf) (Table 1). Primers and probes were synthesized by standard phosphoramidite chemistry techniques at Qingke biotechnology Co. ltd (Beijing, China). TaqMan probes were labeled with the molecule 6-carboxy-fluroscein (FAM) at the 5’ end, and with the quencher Blackhole Quencher 1 (BHQ1) at the 3’ end. Optimal concentrations of the primers and probes were determined by cross-titration of serial two-fold dilutions of each primer/probe against a constant amount of purified RNA of SARS-CoV-2.

**Table 1.**
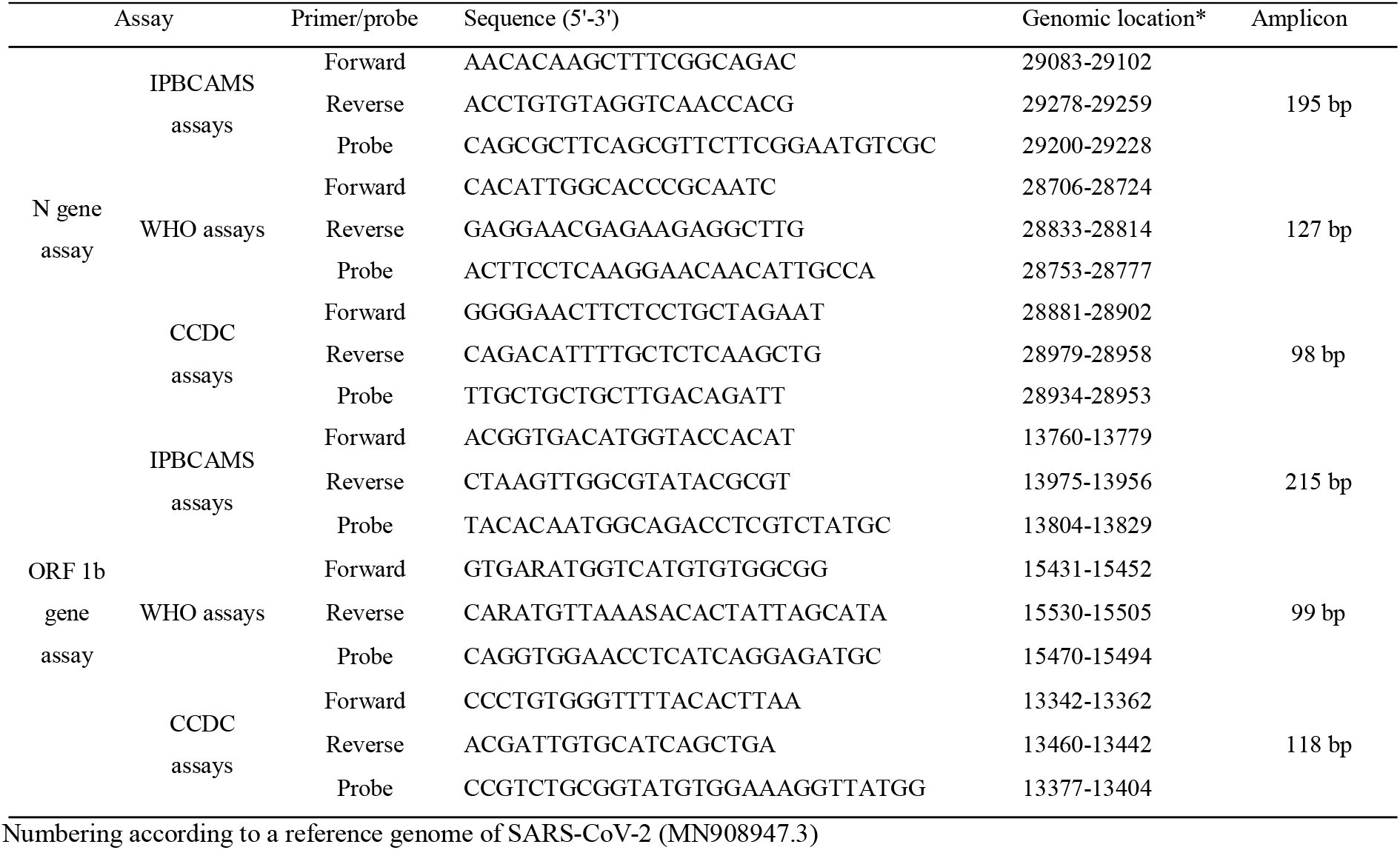
Primers and probes of the three qRT-PCR assays.

### TaqMan real-time RT-PCR assay

The TaqMan real-time RT-PCR assays were performed by using TaqMan Fast Virus 1-Step Master Mix (Thermo Fisher Scientific, MA, USA). Each 20 μl reaction mix contained 5 μl of 4×Fast Virus 1-Step Master Mix, 0.2 μl of 50 μM probe, 0.2 μl each of 50 μM forward and reverse primers, 12.4 μl of nuclease-free water, and 2 μl of nucleic acid extract. Amplifications were carried out in 96-well plates by using Bio-Rad instrument (Bio-Rad CFX96, CA, USA). Thermo-cycling conditions are as follows: 15 min at 50□ for reverse transcription, 4 min at 95□ for pre-denaturation, followed by 45 cycles of 15 sec at 95 and 45 sec at 60□. Fluorescence measurements were taken at 60□ of each cycle. The threshold cycle (Ct) value was determined by the point at which fluorescence exceeded a threshold limit set at the mean plus 10 stand deviations above the baseline. A result was considered positive if two or more of the SARS-CoV-2 genome targets exhibited positive results (Ct ≤ 35). A result of 35 ≤ Ct ≤ 40 was considered suspected and a repeat test was performed for result confirmation.

### Preparation of RNA transcripts

RNA transcripts for N gene and ORF 1b of SARS-CoV-2 were prepared with a plasmid pEasy-T1 (TransGen Biotech, Beijing, China) with T7 promoter before the multiple cloning sites. The plasmids inserted with viral gene regions of N and Orf1b were linearized with the restriction enzyme, BamHI, and transcribed *in-vitro* by using RiboMAX^™^ Large Scale RNA Production Systems (Promega, WI, USA), respectively. The concentrations of the RNA transcripts were determined by using NanoDrop (Thermo Fisher Scientific, CA, USA).

## Results

### Comparison of the sensitivities, reproducibility and linear detection ranges of the three qRT-PCR assays

To determine the sensitivity of the three qRT-PCR assays, we measured the limit of detection (LOD) for each assay by using RNA transcript of the corresponding gene in ten-fold dilution as template (RNA transcript alone). A LOD of 10 genomic copies per reaction was observed for both the N gene assay and the ORF 1b gene assay of all the three qRT-PCR assays, although the Ct values for N gene assay of WHO assays and ORF 1b gene assay of CCDC assays were higher than 35 cycles (Table 2).

**Table 2.** Reproducibility (Coefficient of Variation, %) of the three qRT-PCR assays.

| Assay |  |  | Copy number of RNA transcript |  |  |  |  |  |
| --- | --- | --- | --- | --- | --- | --- | --- | --- |
| | | | $1 \times 10^6$ | $1 \times 10^5$ | $1 \times 10^4$ | $1 \times 10^3$ | $1 \times 10^2$ | $1 \times 10^1$ |
| N gene assay | IPBCAMS assays | Intra-assay | 0.52* | 1.33 | 0.37 | 0.46 | 0.20 | 1.25 |
|  |  | Inter-assay | 1.06 | 2.45 | 1.49 | 1.32 | 1.37 | 1.45 |
|  | WHO assays | Intra-assay | 1.08 | 1.19 | 1.12 | 0.87 | 0.46 | 5.09 |
|  |  | Inter-assay | 7.59 | 2.94 | 2.78 | 6.60 | 0.96 | 3.77 |
|  | CCDC assays | Intra-assay | 0.52 | 0.54 | 0.27 | 0.74 | 0.41 | 1.97 |
|  |  | Inter-assay | 1.56 | 1.20 | 5.51 | 1.00 | 1.40 | 2.89 |
| ORF 1b gene assay | IPBCAMS assays | Intra-assay | 0.73 | 0.26 | 1.10 | 1.30 | 4.45 | 3.36 |
|  |  | Inter-assay | 4.66 | 3.85 | 2.77 | 2.17 | 5.12 | 3.50 |
|  | WHO assays | Intra-assay | 0.57 | 0.47 | 0.88 | 0.41 | 0.29 | 1.76 |
|  |  | Inter-assay | 1.57 | 0.30 | 0.87 | 0.69 | 0.55 | 1.23 |
|  | CCDC assays | Intra-assay | 1.66 | 0.78 | 0.71 | 0.92 | 2.45 | 6.52 |
|  |  | Inter-assay | 0.52 | 0.54 | 0.27 | 0.74 | 0.41 | 1.97 |

The coefficient of variation was calculated by standard deviation of the Ct values of a RNA dilution divided by the mean Ct values of the same RNA dilution.

**Table 3.**
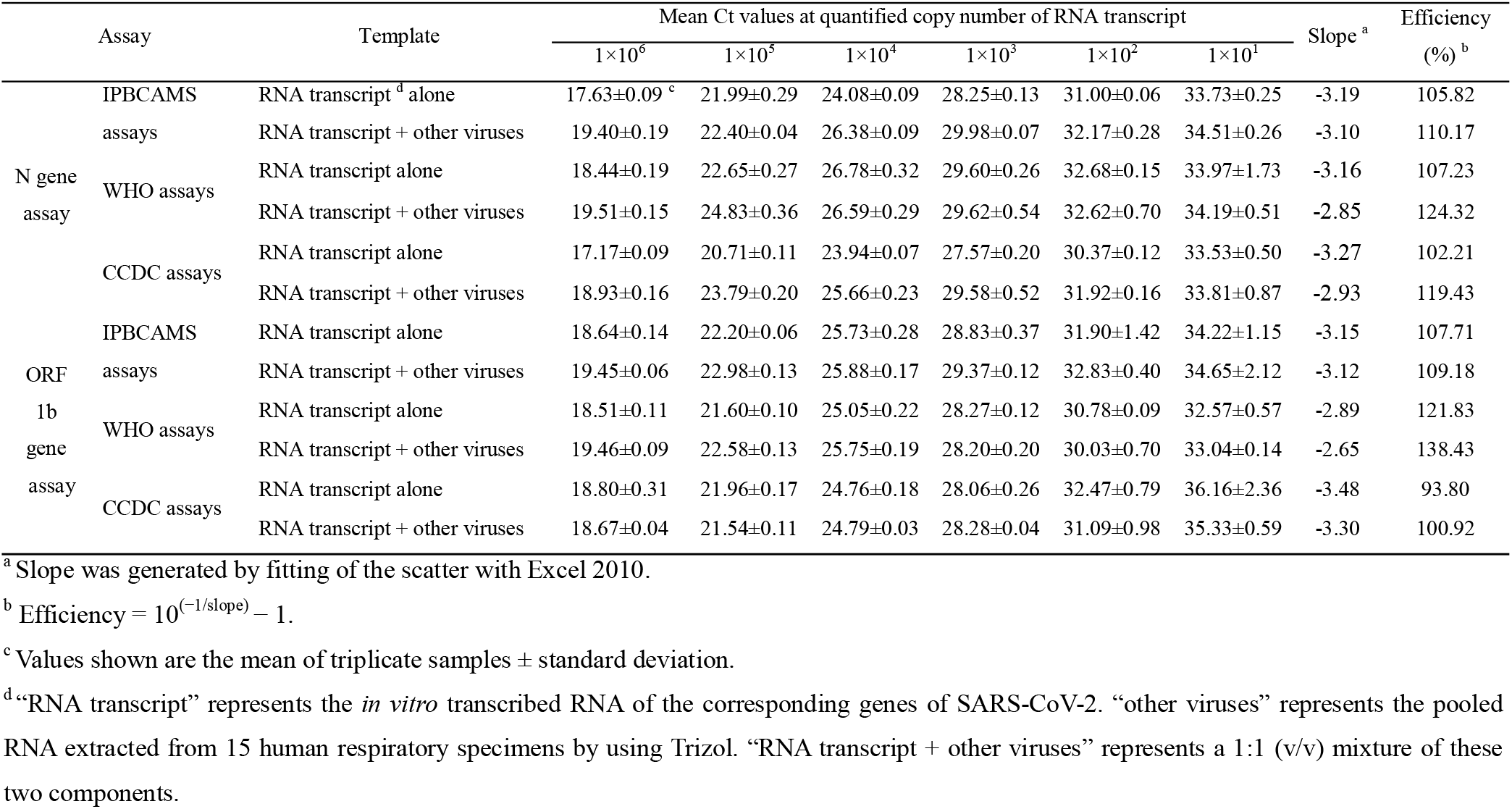
Efficiency of the three qRT-PCR assays.

The linear detection ranges of the three qRT-PCR assays were determined by using a ten-fold dilution of the RNA transcript as template. It showed that the Ct values increased with the RNA transcript from 10^6^ to 10^1^ copies in the reaction in all of the three qRT-PCR assays (Table 2). Strong linear correlations were observed between Ct values and quantity of RNA transcripts with r^2^=0.9926, 0.9750, 0.9987 in the N gene assay, and r^2^=0.9953, 0.9897, 0.9941 in the ORF 1b assay of IPBCAMS assays, WHO assays, and CCDC assays, respectively. These results suggested that all of the three qRT-PCR assays exhibited linear detection ranges from 10^6^ to 10^1^ copies per reaction, while the WHO assays showed lower coefficient of linear correlation.

The reproducibility of the three qRT-PCR assays was assessed by measuring coefficient of variation (CV) of mean Ct values in the intra- and inter- assay. For the N gene assay, the CVs of mean Ct values from 10^6^ to 10^1^ copies of RNA transcript per reaction were 0.20%-1.33%, 0.46%-5.09%, 0.27%-1.97% in intra-assay, and 1.06%-2.45%, 0.96%-7.59%, 1.00%-5.51% in inter-assay of IPBCAMS assay, WHO assay, and CCDC assay, respectively. For the ORF 1b gene assay, the CVs of mean Ct values were 0.26%-4.45%, 0.29%-1.76%, 0.71%-6.52% in intra-assay, and 2.17%-5.12%, 0.30-1.57%, 2.63%-4.34% in inter-assay of IPBCAMS assays, WHO assays, and CCDC assays, respectively.

Because co-infections of respiratory viruses are common, we prepared a (v:v=1:1) mixture of the RNA transcript and a pooled total nucleic acid extract from respiratory specimens (RNA transcript + other extract) as template, to evaluate the effect of co-existed viral nucleic acids on the performance of the assays. No effect of the co-existed other viral nucleic acids on the LOD and the linear detection range was observed, although higher Ct values were generated than those of RNA transcript alone as template in all of the three qRT-PCR assays. However, the co-existed other viral nucleic acids put some effect on the efficiencies of the three qRT-PCR assays.

For the N gene assays, the efficiencies were moved from 105.82%, 107.23%, 102.21% to 110.17%, 124.32%, 119.43% in IPBCAMS assays, WHO assays, CCDC assays, respectively. For the ORF 1b assays, the efficiencies were moved from 107.71%, 121.83%, 93.80% to 109.18%, 138.43%, 100.92% in IPBCAMS assays, WHO assays, CCDC assays, respectively.

### Comparison of the specificities of the three qRT-PCR assays

To evaluate the potential cross-reactions with other human respiratory viruses, the three qRT-PCR assays were examined by using human respiratory samples as templates, which were positive for human coronaviruses (OC43, NL63, 229E, or HKU1), or Influenza viruses (A or B), or respiratory syncytial virus, or parainfluenza virus (1-4), or human metapneumovirus, or rhinovirus, or adenovirus, or bocavirus. No cross reaction was observed in all of the three qRT-PCR assays (data not shown), suggesting high specificity of the three qRT-PCR assays in detecting SARS-CoV-2.

### Assay evaluation with clinical specimens

The three qRT-PCR assays were evaluated with 25 clinical specimens (including 13 throat swabs and 12 sputum) from 25 suspected COVID-19 patients. SARS-CoV-2 was detected from 92% (23/25), 60% (15/25), 100% (25/25) by the N gene assay, and from 64% (16/25), 48% (12/25), 20% (5/25) of all enrolled clinical specimens by the ORF 1b gene assay in IPBCAMS assays, WHO assays, CCDC assays, respectively (Table 4). With respect to the sputum, SARS-CoV-2 was detected from 100% (12/12), 75% (8/12), 100% (12/12) of specimens by the N gene assay, and from 100% (12/12), 75% (8/12), 41.7% (5/12) of specimens by the ORF 1b gene assay in in IPBCAMS assays, WHO assays, CCDC assays, respectively. About the throat swabs, SARS-CoV-2 was detected from 84.6% (11/13), 53.8% (7/13), 100% (12/12) of specimens by the N gene assay, and from 30.8% (4/13), 30.8% (4/13), 0% (0/13) of specimens by the ORF 1b gene assay in in IPBCAMS assays, WHO assays, CCDC assays, respectively. These results demonstrated that the N gene assay performed better than the corresponding ORF 1b gene assay of all the three qRT-PCR assays, the N gene assay in CCDC assays and ORF 1b gene assay in IPBCAMS assays performed better than the other assays.

**Table 4.** Evaluation of the three qRT-PCR assays with clinical specimens.

| Specimen ID | Specimen type | N gene assay |  |  | ORF 1b gene assay |  |  |
| --- | --- | --- | --- | --- | --- | --- | --- |
|  |  | IPBCAMS | WHO | CCDC | IPBCAMS | WHO | CCDC |
| TS98 | Throat swab | 35.79 | NA | 35.42 | NA | NA | NA |
| TS101 | Throat swab | 33.48 | NA | 34.24 | NA | NA | NA |
| TS103 | Throat swab | NA | NA | 34.68 | NA | NA | NA |
| TS105 | Throat swab | 31.5 | 35.76 | 31.64 | NA | NA | NA |
| TS108 | Throat swab | 33.35 | NA | 32.11 | 33.36 | NA | NA |
| TS110 | Throat swab | 29.99 | 31.73 | 29.1 | 33.57 | NA | NA |
| TS165 | Throat swab | 27.34 | 30.46 | 28.14 | 31.06 | 27.84 | NA |
| TS168 | Throat swab | NA | NA | 34.97 | NA | NA | NA |
| TS169 | Throat swab | 33.34 | NA | 34.04 | NA | 34.2 | NA |
| TS187 | Throat swab | 34.5 | 39.2 | 33.03 | NA | NA | NA |
| TS188 | Throat swab | 35.03 | 35.9 | 33.57 | NA | 24.07 | NA |
| TS189 | Throat swab | 31.16 | 35.43 | 31.21 | 34.04 | 30.92 | NA |
| TS190 | Throat swab | 32.84 | 34.02 | 32.56 | NA | NA | NA |
| TY1 | Sputum | 27.35 | 29.44 | 27.6 | 30.98 | 27.33 | NA |
| TY2 | Sputum | 29.38 | 31.26 | 29.06 | 32.32 | 28.72 | NA |
| TY3 | Sputum | 31.85 | NA | 31.3 | 35.84 | NA | NA |
| TY4 | Sputum | 22.99 | 25.57 | 22.08 | 27.42 | 24.12 | 35.99 |
| TY6 | Sputum | 25.51 | 27.52 | 25.58 | 29.03 | 25.58 | 41.54 |
| TY7 | Sputum | 26.9 | 30.21 | 27.4 | 30.05 | 27.3 | 45.26 |
| TY8 | Sputum | 29.21 | 31.87 | 30.06 | 33.65 | 29.84 | NA |
| TY9 | Sputum | 26.29 | 28.45 | 26.34 | 30.69 | 26.03 | 46.34 |
| XT1 | Sputum | 25.74 | 27.26 | 25.3 | 29.82 | 26.34 | 45.9 |
| XT2 | Sputum | 31.57 | NA | 30.95 | 34.19 | NA | NA |
| XT3 | Sputum | 31.14 | NA | 32.02 | 35.02 | NA | NA |
| XT4 | Sputum | 32.67 | NA | 31.71 | 34.26 | NA | NA |
| account (%) of positive |  | 23 (92%) | 15 (60%) | 25 (100%) | 16 (64%) | 12 (48%) | 5(20%) |

## Discussion

Rapid and accurate detection of SARS-CoV-2 represent a fast-growing global demand, which could be met by TaqMan real time RT-PCR (qRT-PCR). However, the current available TaqMan qRT-PCR assays for SARS-CoV-2 are varied in performance, including sensitivity, specificity, reproducibility, linear detection ranges, etc. Due to that relative lower viral load in upper respiratory tract, reliable qRT-PCR assays for the detection of SARS-CoV-2 are required. We thus compared the performance of three currently wide-applied qRT-PCR assays in the detection of SARS-CoV-2. Sensitivity is the primary demand in the detection of respiratory viruses (6). All of the three qRT-PCR assays could provide a LOD of 10 genomic copies per reaction with a detection range from 10^6^-10^1^ genomic copies per reaction. The Ct value at 10 genomic copies per reaction in the ORF 1b gene assay of CCDC assays was higher than 35. These results suggested that most of the three qRT-PCR assays provide high sensitivity and wide linear detection range in detecting SARS-CoV-2, except a relative lower sensitivity observed in the ORF 1b gene assay of CCDC assays.

Specificity is also essential in the detection of SARS-CoV-2, because of common co-infections with other respiratory viruses and high host DNA background in throat swabs (7-9). We evaluated the specificity of the three qRT-PCR assays with respiratory specimens positive for other common respiratory viruses. No cross reaction was observed, demonstrating high specificity of the three qRT-PCR assays in detection of SARS-CoV-2.

We next evaluated the reproducibility of the three qRT-PCR assays by measuring coefficient of variation (CV) of mean Ct values in intra- and inter- assay (10). The N gene assay in IPBCAMS assays and ORF 1b gene assay in WHO assays exhibited a relative better reproducibility with lower intra- and inter- assay CVs, which were not affected by the co-existed nucleic acids of other respiratory viruses.

Efficiency is another key parameter of qRT-PCR, reflecting the binding efficiency of primers & probe to template and the amplification efficiency of the PCR system(11). Most of the qRT-PCR assays provided good efficiency, except an abnormal efficiency of 121.83% observed in the ORF 1b gene assay of WHO assays. An exceptionally high efficiency indicates an increased risk of false positive (12). The co-existed nucleic acids of other respiratory viruses increased the efficiency of all the three qRT-PCR assays, suggesting potential increased risk of cross-reactions between the primers & probe and background nucleic acids.

We finally evaluate the performance of the three qRT-PCR assays with clinical specimens from suspected SARS-CoV-2 infected patients (13). Possibly because of the lower viral load in upper respiratory tract (4), the detection rate of SARS-CoV-2 was lower in throat swabs than in sputum by all of the three assays. Meanwhile, the N gene assay performed better than the corresponding ORF 1b gene assay in all of the three qRT-PCR assays. For the N gene assay, IPBCAMS assays and CCDC assays performed better than WHO assays, both of which could detect SARS-CoV-2 from more than 90% of the suspected specimens. For the ORF1b gene assay, IPBCAMS assays performed better than WHO assays and CCDC assays, with a detection rate of 64%.

In conclusion, we performed methodological evaluations on three widely-applied qRT-PCR assays for the detection of SARS-CoV-2. Although most of the evaluated assays exhibited good sensitivity, specificity, reproducibility and wide linear detection range, performance test with clinical specimens from suspected COVID-19 patients suggested that the N gene assay in IPBCAMS assays and CCDC assays, and the ORF 1b gene assays in IPBCAMS assays were the preferred qRT-PCR assays for accurate detection of SARS-CoV-2.

## Data availability

The original data will be available upon request.

## Acknowledgements

We would like to thank the clinicians who contributed to sample collection and transportation. This study was funded in part by the Project from the Ministry of Science and Technology of China (2020YFC0841200), the National Major Science & Technology Project for Control and Prevention of Major Infectious Diseases of China (2017ZX10103004), the Chinese Academy of Medical Sciences (CAMS) Innovation Fund for Medical Sciences (2020HY320001), the key R&D plan of Shanxi Province (202003D31003/GZ) and the non-profit Central Research Institute Fund of Chinese Academy of Medical Sciences (2019PT310029).

